# Mechanical context defines integrin requirement for maintaining epithelia architecture

**DOI:** 10.1101/2025.09.09.675082

**Authors:** Lourdes Rincón-Ortega, Cecilia H. Fernández-Espartero, Isabel M. Palacios, Acaimo González-Reyes, María D. Martín-Bermudo

## Abstract

Epithelial tissues form cohesive sheets of cells anchored to an underlying basement membrane (BM). They are broadly classified as simple or stratified, each fulfilling distinct physiological roles. While mechanisms driving stratification are increasingly well characterized, those safeguarding simple epithelial architecture remain poorly defined. Here, we address this knowledge gap using the simple follicular epithelium of *Drosophila* as a model. Combining live imaging, quantitative image analysis, manipulation of BM mechanical properties and biophysical measurements, we uncover previously unrecognized integrin-dependent mechanisms esential for preserving simple epithelial integrity. In addition to their known role in orienting cell division, integrins regulate cell reintegration dynamics and modulate surface tension at the cellular level. We further demonstrate that the maintenance of epithelial architecture is governed by mechanisms acting across both cellular and tissue scales. In our model, BM mechanical properties—including stiffness anisotropy—cooperate with integrin-mediated adhesion and tissue geometry to preserve epithelial organization. Given the central role of epithelial disorganization in tumorigenesis, elucidating these mechanical and molecular regulators is critical for understanding epithelial morphogenesis, maintaining tissue homeostasis, and uncovering the early events of cancer progression.

## Introduction

Epithelia, one of the four basic tissue types in the body, consist of continuous sheets of cells standing on a basement membrane (BM). They line internal and external surfaces of all multicellular organisms. Epithelia are broadly classified in two main types, simple, where a single layer of cells is directly attached to the underlying BM, or stratified, which consist of multiple cell layers, with only the basal cells maintaining contact with the BM. Both are essential, serving distinct yet complementary functions. Simple epithelia aid in absorption and secretion, while stratified epithelia primarily protect against physical and chemical damage, and pathogens. Establishing and maintaining both types is therefore critical for organismal health and survival. However, while considerable attention has been given to understanding the formation of stratified epithelia, the mechanisms preserving simple epithelia remain less understood.

Stratified epithelia are believed to originate from simple epithelia. Several studies on epidermal development support a model in which the process of stratification is closely associated with the orientation of cell division within the basal layer ^1–3^. During stratification, epidermal progenitor cells transition from symmetric to asymmetric modes of cell division. In symmetric divisions, the mitotic spindle aligns parallel to both the epithelium and the BM, resulting in the production of two functionally equivalent daughter cells. In contrast, an asymmetric division is characterized by the perpendicular orientation of the mitotic spindle relative to the epithelium and the BM, yielding one self-renewing basal progenitor and one differentiated supra-basal cell ^1, 2^. However, more recent studies have demonstrated that daughter cells resulting from perpendicular divisions within the basal layer may still be reintegrated into the basal compartment following mitosis ^4^. Furthermore, experimental disruption of cell division in the developing skin did not entirely prevent stratification ^4^. These findings indicate that, beyond spindle orientation, alternative mechanisms -such as cellular delamination from the basal to the suprabasal layers-may also have essential functions in epidermal stratification during development. Consequently, the mechanisms governing the transition from simple to stratified epithelia deserve closer attention. Moreover, even less is known about the factors responsible for ensuring not only the configuration but also, critically, the maintenance of simple epithelial architecture.

The mechanical properties of the BM play a critical role in shaping epithelial tissues during development and in maintaining their structural integrity and function. In recent years, a growing body of evidence from studies of epithelial morphogenesis during development strongly supports the notion that spatiotemporal heterogeneities in the BM significantly contribute to tissue shaping and the maintenance of epithelial integrity. Mechanical anisotropy within the BM can arise through spatially and temporally regulated modifications in its degradation, composition, deposition, or the organization of its fibrillar components (reviewed in ^5^. For example, in *Caenorhabditis elegans*, differential BM degradation between the distal tip and the proximal region is critical for gonadal elongation ^6^. Similarly, in *Drosophila*, variations in BM thickness and mechanical stiffness have been shown to regulate anisotropic growth of the wing disc and elongation of the egg chamber, respectively ^7–10^.

Interactions between cells and the BM have also emerged as key determinants in the developmental decision between simple and stratified epithelial architectures, as well as in the preservation of simple epithelial organization. Integrins—heterodimeric transmembrane receptors composed of α and β subunits and the principal mediators of cell adhesion to the BM—have been shown to be essential for proper spindle orientation and regulated cell division during stratification in mouse epidermal development ^1^. Moreover, conditional deletion of β1 integrin (*Itgβ1*) in the developing mouse lung epithelium led to an increased frequency of mitotic spindles oriented perpendicular to the BM, resulting in the transformation of the monolayered epithelium into a multilayered structure ^11^. Similarly, the elimination of integrins from the simple follicular epithelium of the *Drosophila* ovary results in the formation of multilayered epithelial structures ^12^. However, the specific role of integrins in this context remains a subject of debate. Two temporally and spatially distinct functions have been proposed: one implicates integrins in the regulation of mitotic spindle orientation during cell division, paralleling observations in the developing mouse lung and in cell culture ^11, 13^, while the other involves a role in orchestrating cellular rearrangements during the early stages of follicular epithelium development ^14, 15^. Collectively, these findings support the notion that integrins are generally involved in the formation and maintenance of simple epithelial architecture and in preventing epithelial stratification. Nonetheless, the precise mechanisms through which integrins exert these functions remain incompletely understood.

In summary, a substantial body of research conducted in recent years across a range of organisms and model systems has demonstrated that the precise regulation of epithelial tissue architecture, as well as its subsequent maintenance, is governed by mechanisms operating at both cellular and tissue scales. Nevertheless, key factors involved in these processes—such as cell–BM interactions and the mechanical properties of the BM—have often been examined in isolation, despite compelling evidence suggesting that they act in an interdependent manner during epithelial morphogenesis and homeostasis.

The follicular epithelium (FE) of the adult *Drosophila* ovary offers a powerful model to investigate how cell-BM interactions, BM mechanical anisotropy and tissue geometry coordinate to form and maintain simple epithelia. The ovary consists of tubular units known as ovarioles ^16^. Each ovariole contains a germarium at its anterior end and progressively maturing egg chambers towards the posterior end. Each egg chamber comprises a 16-cell germline cyst encased by a monolayer of somatic follicle cells (FCs), known as the follicular epithelium (FE) ^17^. The apical side of FCs faces the germline cells, while their basal surface contacts a BM, which encapsulates the egg chamber ^18^, Fig.1A). Oogenesis spans approximately one week progressing through 14 developmental stages (S1 to S14) ^17^. Upon budding from the germarium, egg chambers are surrounded by approximately 80 proliferating FCs, which switch to endoreplication at S6 ^19^. By S5, FCs located at the anterior and posterior poles of the egg chamber become distinct from central “mainbody” FCs through patterned signalling ^20^. Concomitantly, initially spherical, egg chambers begin to elongate along its anterior–posterior (AP) axis, adopting the elliptical shape of mature eggs ^21^. Throughout this process, FE simple epithelial architecture is maintained by polarity regulators, the cytoskeleton and integrins ^12, 22–25^. Notably, epithelial integrity is most vulnerable at the poles, where loss of polarity or integrins leads to multilayering ^12, 22, 23^. However, the mechanistic basis for this regional susceptibility remains poorly understood.

**Figure 1.**
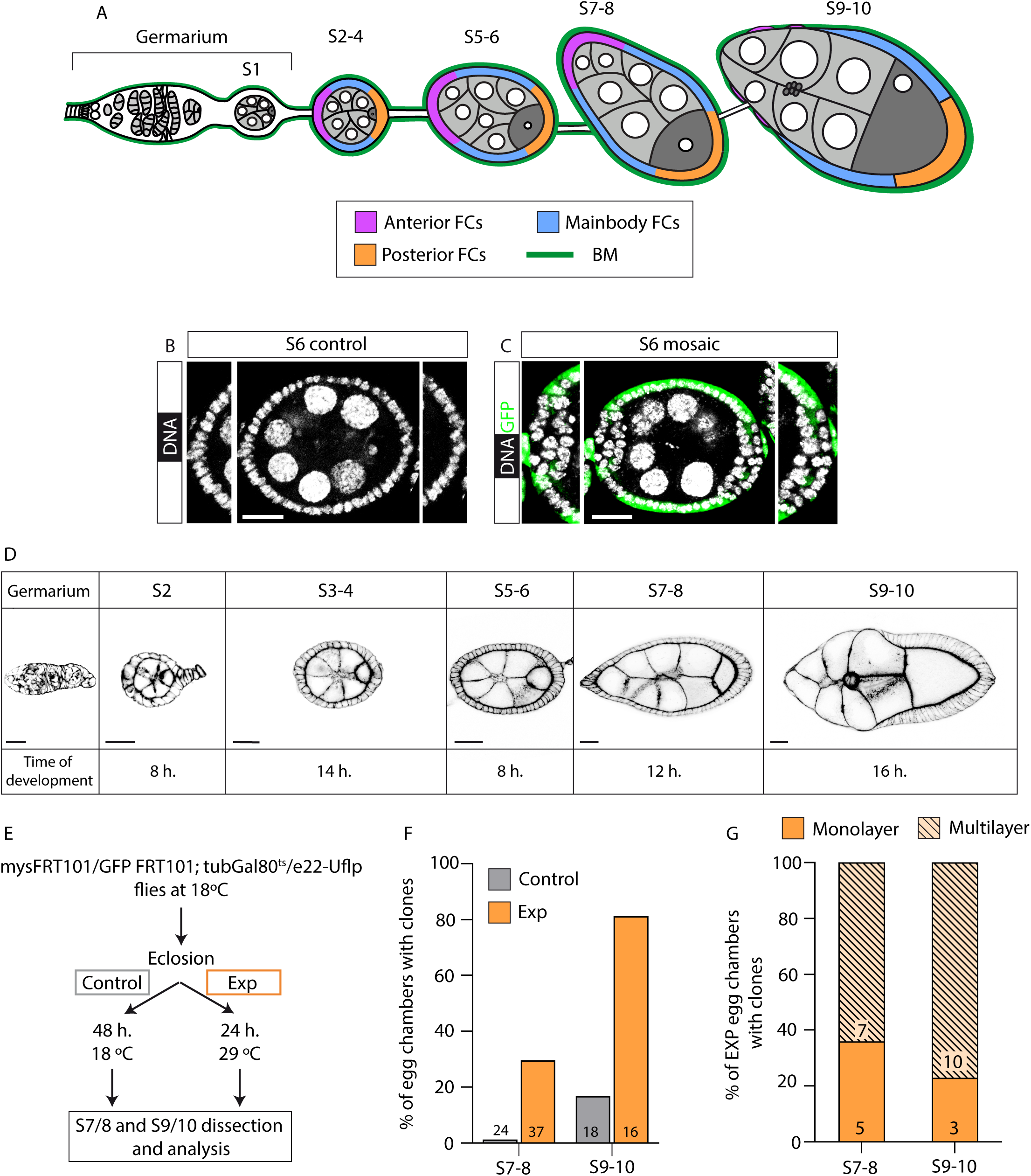
Generation of integrin mutant FCs after the germarium results in multilayering. (A) Schematic diagram of a *Drosophila* ovariole showing the germarium and stages 1 to 10 (S1-S10) egg chambers. Follicle stem cells (FSCs); follicle cells (FCs); stalk cells (StCs); basement membrane (BM). (B) Control egg chamber stained with the nuclei marker TO-PRO-3 (white). (C) Mosaic egg chamber containing *mys* mutant follicle cell clones in the anterior and posterior terminal domains. Mutant FCs are marked by the absence of GFP (anti-GFP in green); TO-PRO-3 (white). (D) A control ovariole visualized with the F-actin marker rhodamine-phalloidin (black), illustrating the germarium and S1 to S10 egg chambers. Developmental timing (in hours) is indicated for each stage. The region shaded in orange denote the time of clonal induction. (E) Schematic of the heat shock regime and clonal analysis timeline. (F) Quantification of the frequency of egg chambers harbouring *mys* mutant follicle cell clones. (G) Quantification of the multilayering phenotype in mosaic egg chambers carrying *mys* mutant follicle cell clones. Scale bars 20 μm (B, C, D).

In this study, we employed the *Drosophila* FE as a model system to investigate at a tissue level the contributions of cell–BM interactions and BM mechanical properties, and, critically, their coordinated interplay in the morphogenesis and homeostasis of simple epithelia. Through the application of real-time imaging, biophysical approaches—including laser ablation and atomic force microscopy—and quantitative image analysis, we demonstrate that integrin-mediated cell–BM interactions are essential for both the formation and maintenance of the FE. These interactions regulate key processes, such as the orientation of the division plane, cell reintegration following non-planar divisions, and the modulation of cortical tension. Furthermore, by manipulating BM stiffness, we demonstrate that simple epithelial architecture is ultimately preserved by the integrated action of cell–BM adhesion, spatial mechanical heterogeneities within the BM, and, likely, tissue geometry.

## Results

### Integrins are required for the formation and maintenance of the FE monolayered epithelium

The *Drosophila* genome encodes two β integrin subunits—βPS and βν—and five α subunits, designated αPS1 through αPS5 ^26, 27^. Of these, βPS, encoded by the *myospheroid* (*mys*) gene, is the only β integrin subunit expressed in the ovary ^12, 26, 27^.

The role of integrins in the formation and maintenance of the FE remains contested. Early studies showed that loss of integrin function in clones of FCs led to multilayering at the egg chamber poles and increased spindle misorientation, suggesting a role for integrins in epithelial organization via spindle alignment ^12^, Fig. 1B-C). In contrast, a subsequent study employing temporally controlled, global reduction of integrin levels in all FCs found that while integrin depletion in the germarium induced multilayering at the poles, its elimination after this stage had no discernible effect ^14^. This study also showed severe disorganization of the FE following integrin loss, including the detachment and internalization of pre-follicle cells from the BM, proposed to disrupt the monolayered architecture. Further research revealed that spindle misorientation alone does not disrupt epithelial integrity, as misplaced cells can reintegrate ^28^. Collectively, these findings led to the proposal that the primary function of integrins in the FE is not the regulation of spindle orientation, but rather the maintenance of cellular organization and morphology—a role that appears to be critical only within the germarium ^15^.

Two key methodological differences distinguish our study from that of Lovegrove et al. (2019) with respect to the elimination of integrin function in FCs. First, our approach abrogated integrin function throughout FC development, including both early and late stages, rather than restricting integrin loss to post-germarial stages. Second, while Lovegrove et al. reduced integrin levels in all FCs, by expressing a *mys*-targeting RNAi, we employed mitotic recombination to generate mosaic FE containing a mixture of control and *mys* null mutant FCs. To determine whether these methodological differences could account for the discrepancies in phenotypic outcomes reported, we used a system that enables temporal control of *mys* mutant clone induction in a temperature-dependent manner (Fig. 1D). In this system, clonal induction is suppressed at 18 °C (control condition) and activated at 29 °C (experimental condition). By shifting flies from 18 °C to 29 °C, at defined intervals post-eclosion, we were able to control the developmental stage at which *mys* clones were induced (see Materials and Methods). Based on the timing of egg chamber progression ^29^, Fig.1D, E), stages 7/8 and 9/10 analyzed 24 hours after temperature shift would be expected to harbor *mys* mutant progeny corresponding to clonal induction at stages 3/4 and 5/6, respectively (see Materials and Methods, Fig.1D, E). Consistent with this, control flies maintained at 18 °C exhibited *mys* clones at stages 7/8 and 9/10 in only 1,2% and 16,7% of egg chambers, respectively (Fig.1F). In contrast, flies shifted to 29 °C showed a substantially higher incidence of *mys* clones at the same stages—29,7% and 81,3%, respectively (Fig. 1F). Among these, 64% and 77% exhibited multilayering (Fig.1G), a phenotype never observed in control clones. These findings demonstrate that the loss of integrin function in a subset of FCs, even after they exit the germarium, is sufficient to disrupt epithelial architecture and induce multilayer formation.

Having established that integrins are not only essential for maintaining cellular organization and morphology in the germarium ^15^, but also for preserving the monolayered architecture of the FE, we next sought to investigate the mechanisms through which integrins fulfill this role.

### Integrins regulate FC division plane orientation

The role of integrins in the regulation of spindle orientation in the FE remains a subject of debate. We previously reported that integrin mutant FCs exhibit a higher frequency of spindle orientations perpendicular to the plane of the FE compared to their wild-type counterparts ^12^. In contrast, a later study found no significant difference in spindle orientation between integrin mutant and control FCs when cells remained within the epithelial layer, with increased variability observed only in multilayered regions, most likely as a secondary consequence of epithelia disorganization ^14^. To try to clarify this discrepancy, we performed live imaging to monitor cell divisions in both control and *mys* mutant FCs, during stages 3 to 6 of egg chamber development.

Previous *in vivo* studies demonstrated that FCs can divide either parallel to the FE plane or at an angle ^28^. To assess whether integrin loss alters this behavior, we analyzed and categorized FC divisions based on spindle angle and positioning of the daughter cells. We define two types of divisions, planar and non-planar. Planar divisions were characterized by spindle orientation at an angle of 0–25° relative to the epithelial plane, resulting in both daughter cells remaining within the monolayer (Fig. 2A, B; Movie S1). In contrast, non-planar divisions were defined by spindle angles between 35–90°, wherein one daughter cell was displaced outside the monolayer (Fig. 2A, C; Movie S2). To explore the basis of regional specificity in multilayer formation, we analyzed FC divisions in both polar and mainbody regions of the FE.

**Figure 2.**
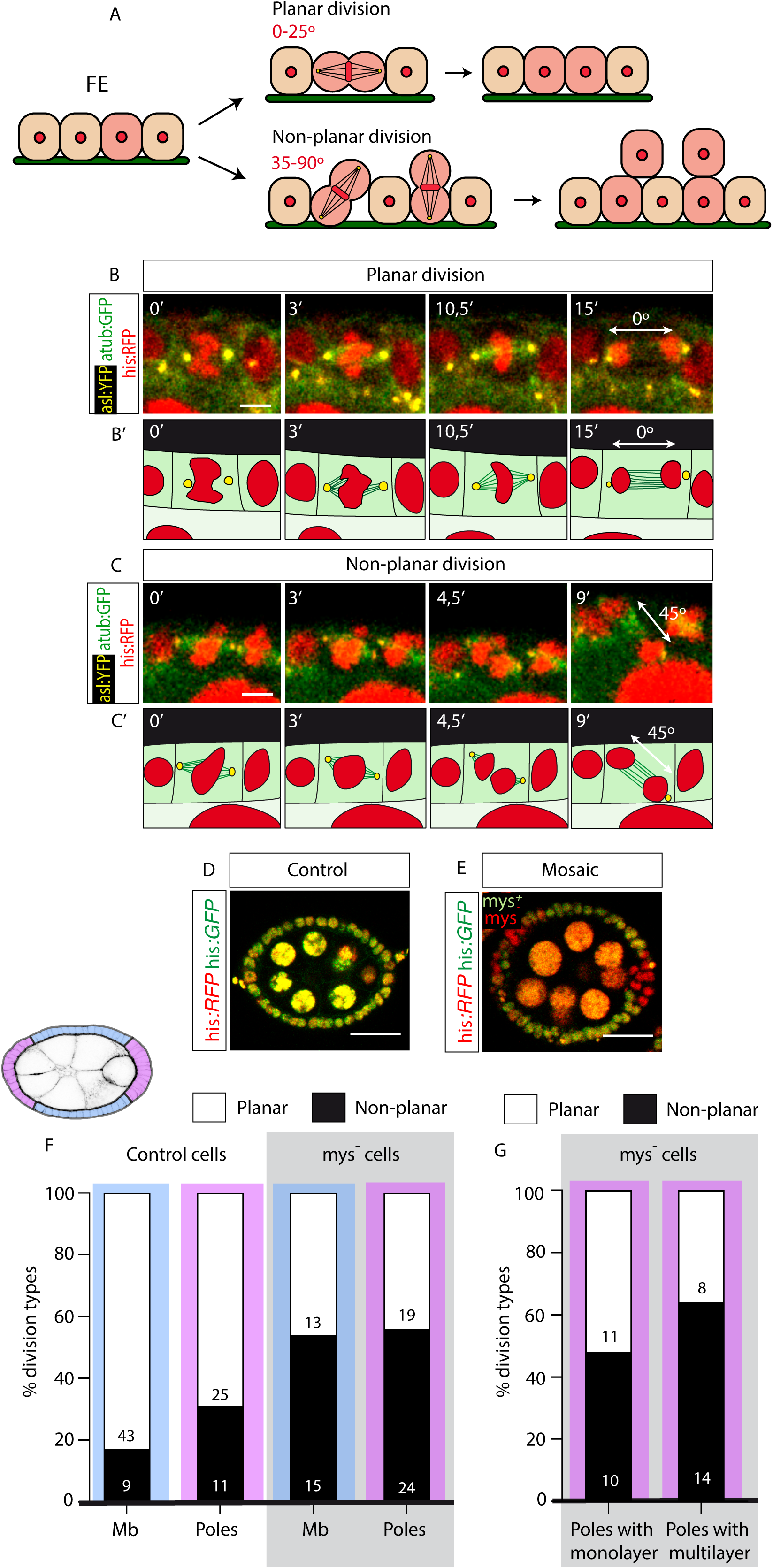
Regional and integrin-dependent bias towards non-planar versus planar divisions in FCs. (A) Schematic representation of planar and non-planar divisions in the follicular epithelium (FE). (B, C) Confocal images of a live S6 egg chamber expressing histone-RFP (his:RFP in red), αtubuline-GFP (αtub:GFP in green) and asterless YFP (asl:YFP in yellow), illustrating examples of a planar division (B) and a non-planar division (C) at the mainbody. (B’, C’) Schematics corresponding to the divisions shown in B and C, respectively, highlighting spindle orientation. (D, E) S5 control (D) and S6 mosaic egg chambers containing *mys* mutant follicle cell clones in the anterior and posterior terminal domains (E), expressing his:RFP (red) and his:GFP (green). Mutant FCs are marked by the absence of his:GFP (green). (F) Quantification of planar versus non-planar division frequencies in control and mosaic egg chambers harbouring *mys* mutant clones, separated by region: mainbody (Mb) and poles. (G) Quantification of division orientation in mosaic egg chambers carrying *mys* mutant follicle cell clones with or without multilayering. Scale bars 3 μm (B, C) and 20 μm (D, E).

Notably, in control egg chambers (Fig. 2D), non-planar divisions occurred more frequently at the poles (30,5%, n = 36) compared to the mainbody (17.3%, n = 52), indicating an intrinsic regional difference in division orientation within the wild-type FE (Fig. 2F). In mosaic egg chambers containing *mys* mutant clones (Fig. 2E), the proportion of non-planar divisions among mutant FCs in the mainbody was significantly increased compared to controls (53.5%, n = 28; Fig. 2F). Similarly, the frequency of non-planar divisions at the poles was significantly higher in mosaic egg chambers than in controls (55.8%, n=43). As noted above, integrin loss does not always lead to multilayering. Interestingly, we observed that the increase in non-planar divisions in mosaic egg chambers was more pronounced in those displaying the multilayer phenotype compared to those maintaining a monolayer (63,6%, n=22 versus 47,6%, n=21, Fig.2G).

These findings suggest that while a basal level of non-planar divisions is tolerated without compromising monolayer integrity, a higher frequency—particularly at the poles— correlates with multilayer formation, implicating defects in spindle orientation as a contributing factor. Moreover, they further support a role for integrins in regulating the plane of cell division.

### Integrins facilitate cell reintegration following non-planar division

Previous work has shown that divisions occurring at an angle did not disrupt epithelial architecture, as displaced daughter cells were capable of reintegrating into the monolayer ^28^. To investigate whether integrins play a role in this reintegration process, we conducted live imaging to monitor and characterize reintegration dynamics in both control and mosaic epithelia.

As anticipated, no significant difference in reintegration time was observed between control cells (n=70) and *mys* mutant cells (n=19) undergoing planar divisions (Sup. Fig. 1; Movies S3, S4).

In the context of non-planar divisions, reintegration times varied by region in control egg chambers (Fig. 3). FCs dividing at the poles required significantly more time to reintegrate into the monolayer (mean 40.3 min; n=23) compared to those in the mainbody (mean 20.3 min; n=13) (Fig. 3A, C; Movies S5, S6). In mosaic egg chambers, *mys* mutant FCs in the mainbody exhibited prolonged reintegration times (mean 61.5 min, n = 23) relative to controls (Fig.3B, C, Movie S7). Similarly, *mys* mutant FCs at the poles required significantly more time (mean 80,5 min; n=30) to reintegrate than both control FCs at the poles and mutant cells in the mainbody (Fig.3B, C, Movie S8). The extended reintegration time of integrin mutant cells at the poles may help explain the regional specificity of the multilayering phenotype. In this context, a longer residence time outside the epithelial layer may increase the likelihood of displaced cells undergoing division in the ectopic layer, thereby contributing to multilayer formation. Consistent with this, integrin mutant cells located in ectopic layers exhibited a significantly longer reintegration time (mean 96.6 min., n=15, Fig.3D, Movie S9) compared to mutant cells within a monolayer (mean 64,4 min., n = 15, Fig.3D, Movie S8).

**Figure 3.**
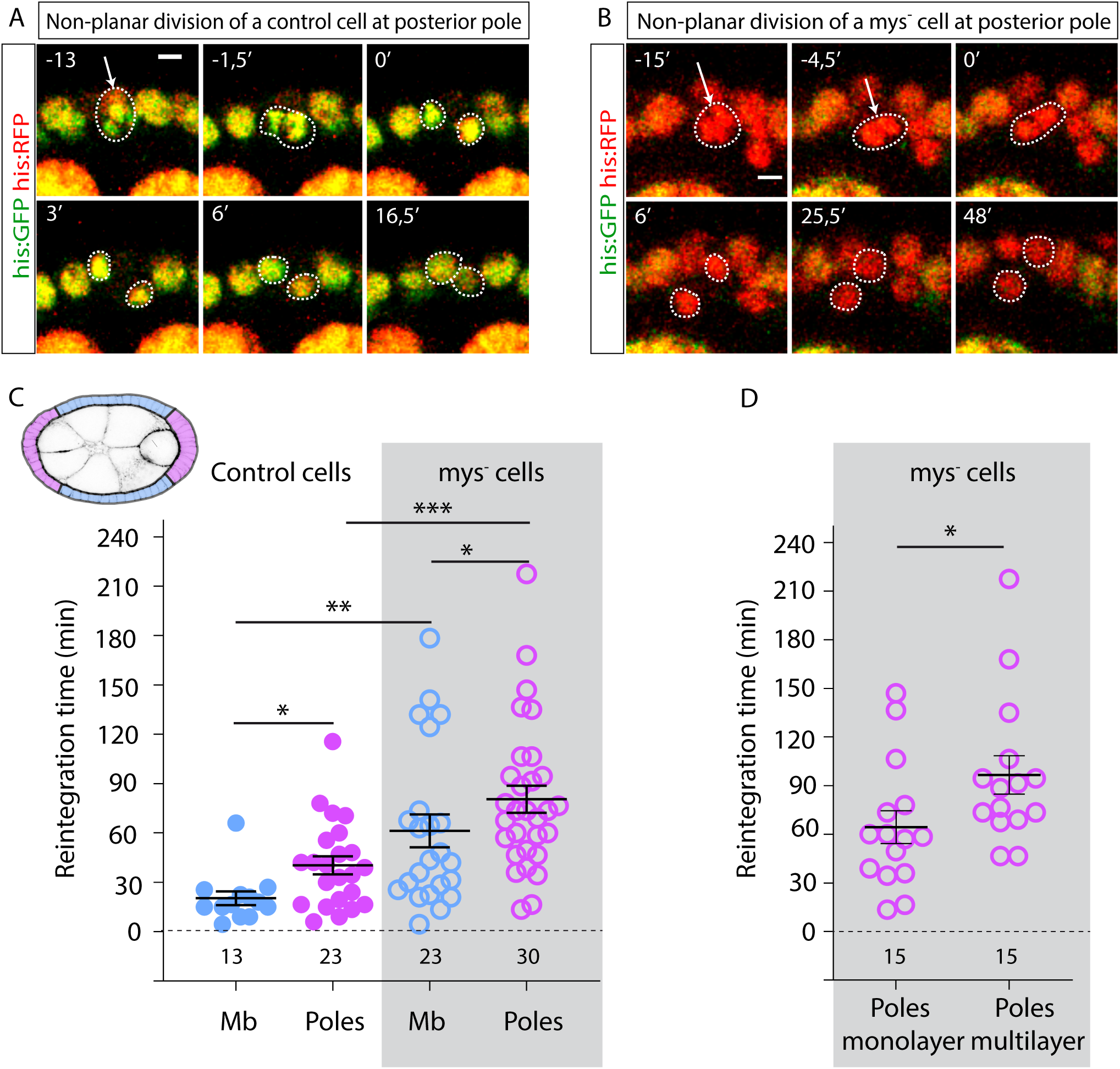
Regional and integrin-dependent bias towards longer reintegration times following non-planar divisions. (A, B) Confocal images of live S5 egg chambers expressing histone-RFP (his:RFP, red) and histone-GFP (his:GFP, green), showing examples of cell reintegration following a non-planar division at the poles in a control (A) and a mosaic egg chambers containing *mys* mutant follicle cell clones (B). Mutant FCs are identified by the absence of his:GFP (green). (C) Quantification of reintegration times in control and mosaic egg chambers harbouring *mys* mutant clones, separated by region: mainbody (Mb) and poles. Schematic in the top left corner illustrates a S6 egg chamber, with the mainbody region shaded in pale blue and the poles in pink. (D) Quantification of reintegration times in *mys* mutant FCs within mosaic egg chambers, comparing clones with and without multilayering. The statistical significance of differences was assessed with a t-test, ***, ** and * P value < 0.0001, <0.001 and <0.05, respectively. Error bars indicate standard error of the mean (s. e). Scale bars 3 μm (A, B).

These results indicate that integrins are essential not only for controlling division orientation, but also for ensuring efficient reintegration of misplaced FCs, thereby safeguarding epithelial architecture. Moreover, they suggest the existence of a threshold reintegration time beyond which the integrity of the monolayer is compromised.

### Multilayered integrin-deficient cells show increased tension at cell-cell contacts

We have previously shown that loss of integrin function leads to increased surface tension at cell-cell contacts in S9 FCs ^30^. These findings raise the possibility that the role of integrins as modulators of tension at cell-cell contacts may also contribute to the formation of multilayers in the FE. To explore this hypothesis, we quantified membrane tension in stage S5/6 control and *mys* mutant FCs.

Laser ablation of cell-cell contacts is a well-established method to measure tension at junctional membranes ^31^. To investigate whether integrin loss affects membrane tension in stage S5/6 FCs, we performed laser ablation experiments using a UV laser beam to sever plasma membranes and underlying cortical cytoskeleton on the basal side of stage S5/6 control and *mys* FCs (see Materials and Methods, Fig. 4). Membrane behaviour was visualised using Resille-GFP ^32^. Because retraction velocity is affected by cytoplasmic viscosity ^33^, we assumed viscosity to be equivalent in *mys* and control FCs. To minimise potential effects due to an anisotropic distribution of forces in the FE, all cuts were made parallel to the anterior-posterior (AP) axis. To assess potential regional differences, ablations were performed in both the polar and mainbody regions of the FE (Fig. 4A, B). As expected, the severing of the membrane resulted in a relaxation of tension at cell-cell junctions, leading to an increase in the distance between cell vertices on either side of the ablation site (Fig. 4B; Movie S10). In control egg chambers, the initial retraction velocity was comparable between the mainbody (0.43 μm/sec, n=16) and pole regions (0.42 μm/sec, n=27) (Fig. 4C). Similarly, *mys* mutant FCs that were not part of a multilayer displayed no significant differences in vertex displacement compared to controls, with velocities of 0.40 µm/sec in the mainbody (n=16) and 0.42 µm/sec at the poles (n=22). In contrast, *mys* mutant FCs located within multilayered regions exhibited significantly greater vertex displacement over time, with an average retraction velocity of 0.65 µm/sec (n=19) (Fig. 4C).

**Figure 4.**
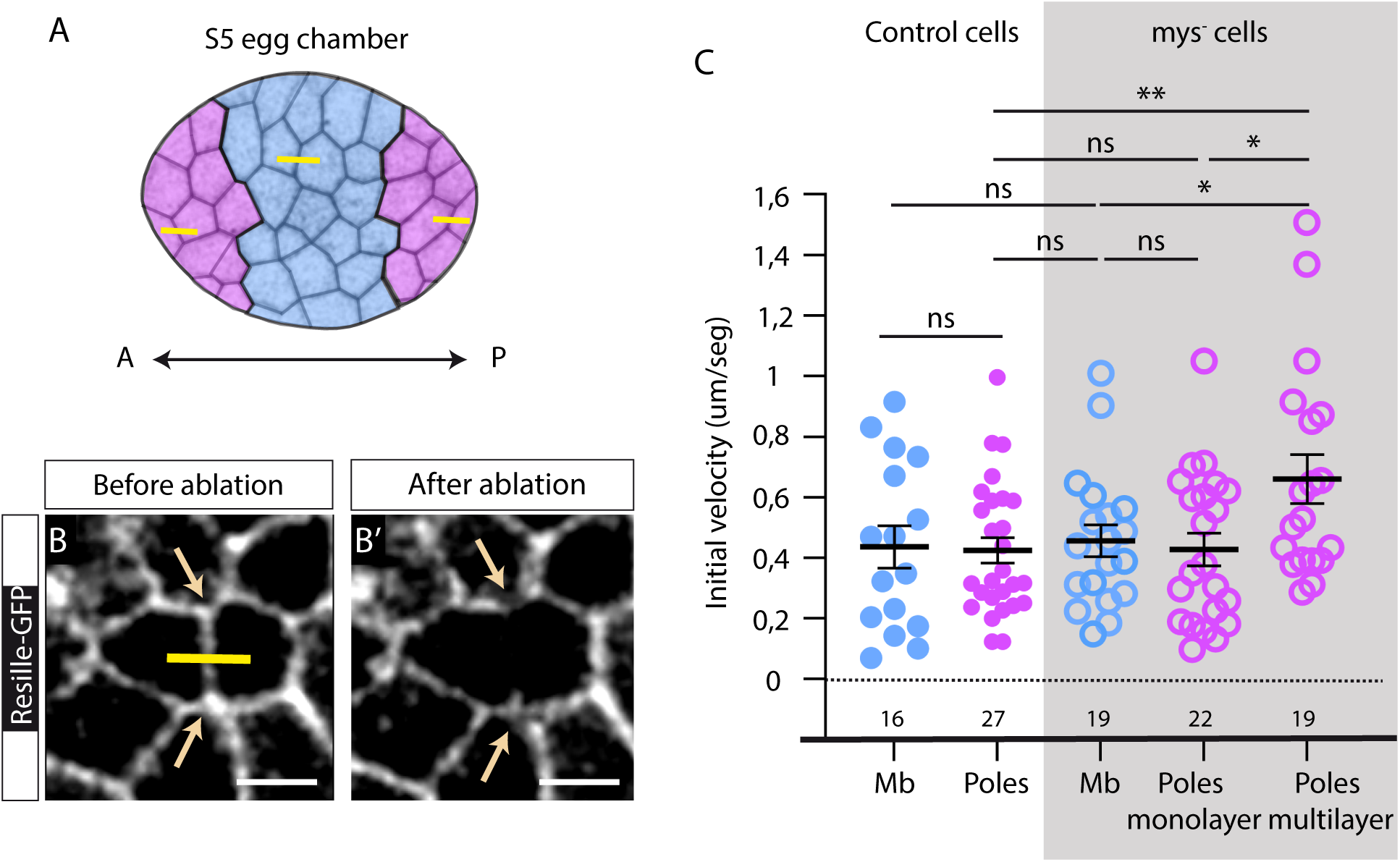
Loss of integrins results in increased membrane tension in FCs located in ectopic cell layers. (A) Schematic of a S5 egg chamber, with the mainbody region shaded in pale blue and the poles in pink. A, anterior; P, posterior. (B) Representative images of life S6 control egg chambers expressing Resille-GFP, before and after laser ablation of individual cell-cell junctions. (C) Quantification of initial velocity of vertex displacement following ablation of the indicated cell junctions. Statistical significance of differences was determined using a t-test, ** and * P value <0.001 and <0.05, respectively; ns not significant. Error bars indicate standard error of the mean (s. e). Scale bar 3 μm (B).

These findings demonstrate that tension at cell-cell contacts is high in integrin mutant FCs that delaminate and remain within multilayered regions. This increase in membrane tension may promote their separation from the layer in contact with the germline and impede their reintegration. In this context, cell proliferation within the multilayered regions could contribute to the generation of additional layers. If this mechanism is operative, one would expect the severity of the multilayering phenotype to progressively increase as egg chamber development advances. Indeed, we found a developmental stage-dependent rise in the proportion of mosaic egg chambers displaying a multilayered phenotype (Sup. Fig.2A-I).

Cells with high interfacial tension have been shown to segregate and sort into distinct groups within a tissue ^34^. Given that integrin mutant follicle cells (FCs) exhibit increased cortical tension relative to their control neighbors, and that substantial cell rearrangements occur during the retraction of main body FCs at S9, these factors may together promote the progressive clustering of integrin mutant cells. In line with this, we find that integrin mutant FCs located at the poles increasingly segregate from neighbouring cells and cluster over time, within ectopic layers (Sup. Fig.2J).

### Spatial heterogeneity in the BM mechanical properties critically governs the formation and maintenance of the simple FE

As mentioned in the Introduction, heterogeneity in the mechanical properties of the BM contributes to the precise regulation of tissue architecture during development and its subsequent maintenance (reviewed in ^5^. Measurements of BM stiffness in the *Drosophila* egg chambers have revealed a robust mechanical anisotropy in the BM, with the poles being approximately 50% softer than the central regions at stages 3 and 5 ^8^. In our study, we observed a clear asymmetry in both division plane orientation and the reintegration time of FCs between the poles and the mainbody of control egg chambers (Figs. 2 and 3). In addition, we find that loss of integrins leads to multilayer formation specifically at the poles, even though integrin mutant FCs in the mainbody also display increased frequencies of non-planar divisions and prolonged reintegration times compared to controls. This suggests that regional differences in BM stiffness may underlie the observed phenotypic disparities.

To test whether BM stiffness heterogeneity contributes to multilayering in the absence of integrins, we analyzed the behavior of integrin mutant cells in egg chambers with homogenously soft BMs. Integrin downregulation in the FE reduces laminin levels ^35^, which in turn decreases BM stiffness ^8^. Thus, global integrin downregulation may produce a uniform softening of the BM, providing an ideal context in which to assess the role of BM stiffness heterogeneity in driving multilayering in the absence of integrins. To reduce integrin levels across the entire FE, we expressed an RNAi construct targeting *mys* under the control of the *tj*-Gal4 driver (*tj*>*mys*RNAi) and measured BM stiffness (Fig.5A-C). Although previous studies have shown that strong integrin knockdown can result in severe disorganization of the ovariole structure ^14^, thereby complicating the analysis of FE-specific phenotypes, we used an alternative *mys*RNAi line ^36, 37^, that reduces integrin levels without inducing major structural defects (Fig. 5A, B). Expression of this RNAi construct throughout the FE results in rounded egg chambers that fail to elongate, consistent with previously reported observations ^15^. To quantify BM stiffness, we used atomic force microscopy to measure the apparent elastic modulus (K), as a proxy for stiffness ^35, 38^. As expected, *mys*-depleted follicles (n=14 from 6 different females) exhibited significantly reduced K values compared to controls (n=16 from 6 different females, Fig. 5D). Furthermore, the typical anisotropic distribution of BM stiffness observed in wild type egg chambers was lost in *tj*>*mys*RNAi egg chambers (Fig.5D). We next examined division orientation and reintegration dynamics, using the membrane marker Resille-GFP to visualize cell behavior live. As described above, control egg chambers (*tj*-ResilleGFP) displayed region-specific rates of non-planar divisions, with approximately 28,95% in the mainbody and 38,71% at the poles (Fig. 5E, Movies S11, S12). In contrast, experimental egg chambers (*tj*-ResilleGFP>*mys*RNAi) showed elevated frequencies of non-planar divisions in both regions (42,86% in the mainbody and 58,82% at the poles, Fig. 5E). Similarly, analysis of reintegration times revealed regional disparities in controls, averaging 29,12 min. in the mainbody and 42,43 min. at the poles, whereas these differences were abolished in *tj*-ResilleGFP>*mys*RNAi egg chambers. In the latter, reintegration times were prolonged across both regions, averaging 51,87 min. in the mainbody and 65,23 min. at the poles (Fig.5F). Notably, these values remain substantially lower than those observed in *mys* mutant FCs within mosaic egg chambers that retain BM stiffness anisotropy (mean: 96.6 minutes). Of interest, *tj*-ResilleGFP>*mys*RNAi egg chambers exhibited multilayering at the poles only sporadically, occurring in 23,61% (n=38) of the cases analyzed (Fig. 5C), compared to 78,37% (n=29, Sup. Fig. 3G) in mosaic egg chambers. Furthermore, when multilayering did occur, it remained limited in extent, never exceeding two layers. These results may explain the lack of a robust multilayering phenotype upon integrin depletion in all FCs, thereby reconciling discrepancies between our findings and previous reports ^14^.

**Figure 5.**
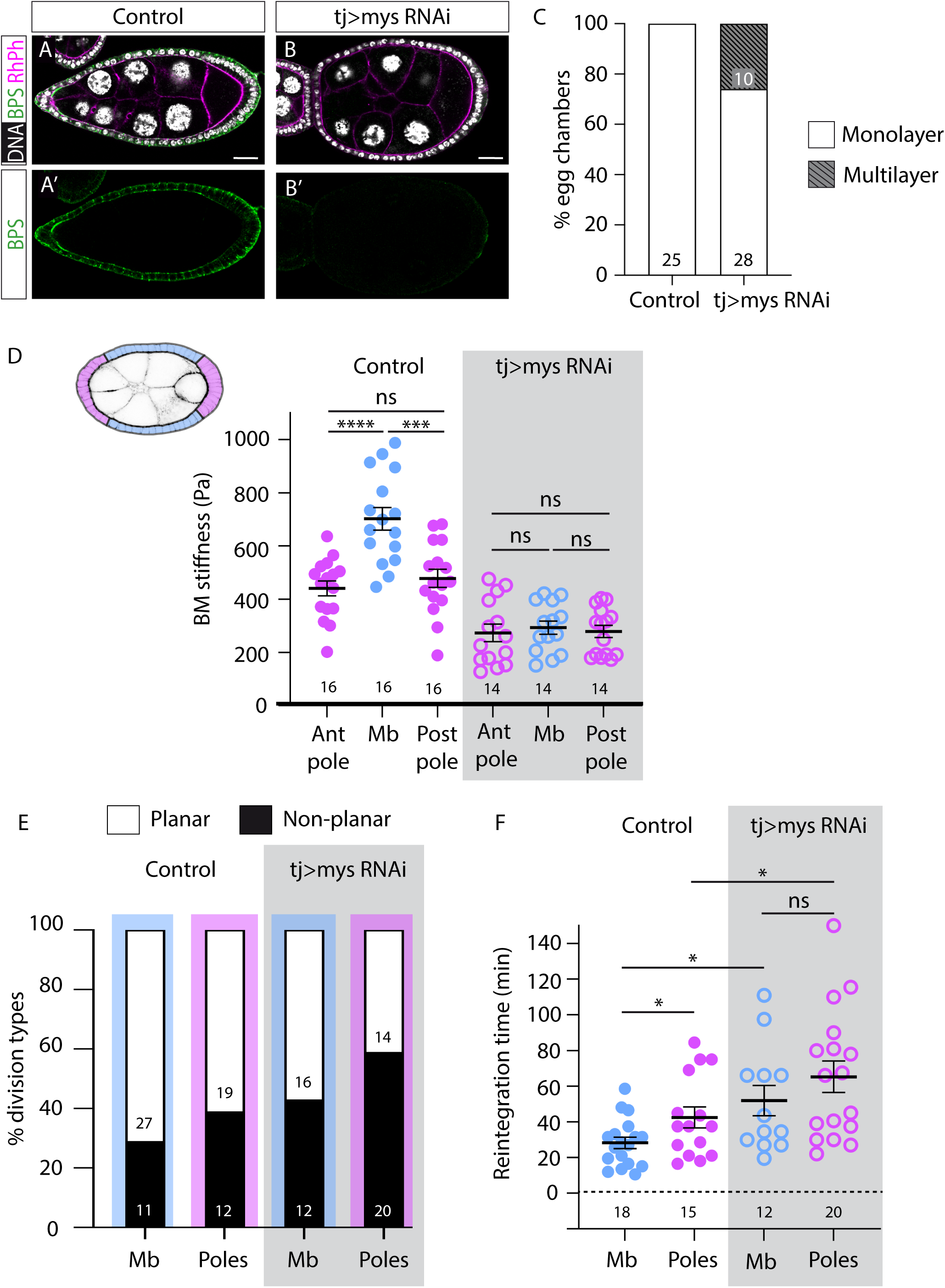
Uniform loss of integrin function in the FE does not induce multilayering. (A, A’) Control and (B, B’) egg chambers with uniform *mys* depletion (*tj>mys* RNAi) stained with the nuclei marker Hoechst (white in A, B) and an antibody against the βPS integrin subunit (green, A, A’, B, B’). (C) Quantification of the multilayering phenotype in control and *tj>mys* RNAi egg chambers. (D) Schematic of a S6 egg chamber, with the mainbody region shaded in pale blue and the poles in pink. (E) Comparison of the apparent elastic modulus *K* in the mainbody (Mb), anterior pole (Ant pole) and posterior pole (Post pole) regions of control and *tj>mys* RNAi egg chambers. (F, G) Quantification of planar versus non-planar division frequencies (F) and reintegration times (G) in control and *tj>mys* RNAi egg chambers, separated by region: mainbody (Mb) and poles. The statistical significance of differences was assessed with a t-test, *** and * P value < 0.0001 and <0.05, respectively.; ns not significant. Error bars indicate standard error of the mean (s. e). Scale bar 20 μm (A, B).

Altogether, these results strongly support a critical role for the spatial heterogeneity in BM mechanical properties in the formation and maintenance of the simple FE. To further test this statement, we generated integrin mutant clones in a context where BM stiffness is homogenous. Egg chambers uniformly depleted of laminins or Collagen IV (Col IV) exhibit homogenously soft BMs ^35^. Thus, we generated *mys* mutant clones in egg chambers depleted of either laminins or ColIV. Laminin levels were reduced in all FCs, using the *tj*-Gal4 driver to express an RNAi construct targeting *Laminin B1* (*LanB1*), which encodes the sole β chain shared by the two Laminin trimers present in *Drosophila* ^35, 39^. To reduce Col IV levels across the FE, we used the *tj*-Gal4 driver to express an RNAi construct targeting *viking* (*vkg*), which encodes *ColIVα2*, one of two Collagen IV subunits in *Drosophila* (*tj*>*vkg*RNAi). Strikingly, reducing either Laminin or ColIV levels substantially rescued the multilayering phenotype. While 78,37% of mosaic egg chambers containing *mys* clones exhibited multilayering (n=29), only 26% of *tj*>*LanB1*RNAi (n=50) and 28,57% of *tj*>*vkg*RNAi egg chambers with *mys* clones (n=56) developed a stratified epithelium (Sup. Fig. 3).

Taken together, these findings demonstrate that the mechanical heterogeneity of the BM contributes to the spatial specificity of integrin-dependent disruptions in a monolayered epithelial architecture. More broadly, our results are consistent with the notion that the preservation of epithelial architecture is regulated by mechanisms acting across both cellular and tissue scales.

## Discussion

Epithelia are continuous cell layers supported by a basement membrane, lining internal and external body surfaces. Simple and stratified epithelia perform essential, distinct functions in the body, yet the mechanisms that maintain simple epithelial architecture remain largely unexplored, compared to those governing stratification. In this study, we employed the monolayered FE of the *Drosophila* ovary as a model system to identify novel regulators of simple epithelial architecture. Our findings reveal that the formation and maintenance of single epithelial layer are governed by multiple factors operating at both cellular and tissue levels.

Previous studies have provided evidence supporting a role for integrins in maintaining simple epithelial architecture and suppressing stratification. Downregulation of *Itgb1* integrin expression appears to be part of the normal stratification program in the human epidermis ^40, 41^ and loss of *Itgb1* leads to abnormal multilayering in both skin and lung tissues ^42^. Our current findings build upon our previous work proposing that integrins are crucial for maintaining the monolayered follicular epithelium (FE) in *Drosophila*, in part by promoting planar rather than non-planar cell divisions ^12^. Consistent with this, integrin-deficient FCs within the monolayer exhibit a significant bias toward non-planar divisions compared to controls. Moreover, mutant cells located in ectopic layers display an even higher frequency of non-planar divisions than their counterparts within the monolayer, which we propose contributes to the reinforcement of multilayering. These observations align with recent *in vivo* studies in mouse skin explants, where continued division of suprabasal cells has similarly been proposed to aid in multilayer formation ^4^. The consistent observation that integrin loss biases FCs towards non-planar divisions reinforces the established role of integrins in mitotic spindle orientation. In cell culture systems, integrin β1 has been proposed to regulate spindle positioning via a ligand-independent mechanosensory complex, comprising FAK, Src, and p130Cas ^13^. Future work should aim to determine whether this pathway is conserved *in vivo* and operates within intact epithelial tissues, and further elucidate the molecular mechanisms by which integrins regulate cell reintegration dynamics.

Although our findings support a role for integrins in controlling spindle orientation and the plane of division ^12, 13^, a compelling aspect of multilayer formation in the *Drosophila* FE is that integrin depletion does not invariably lead to multilayering. This raises the unresolved question of whether disruption of simple epithelial architecture—or its transition to a stratified state—necessarily depends on a shift toward perpendicular division orientation. Although numerous studies in mammalian epithelial tissues, including the skin, hair follicle, tongue and prostate, have highlighted the importance of oriented cell divisions in ensuring proper morphogenesis and structure of mammalian epithelial tissues ^1, 43–45^, other findings challenge this paradigm. Thus, skin explant cultures treated with inhibitors of cell division retain some capacity to stratify and differentiate ^4^. Furthermore, in embryonic mouse kidney, zebrafish neuroepithelial cells, and many *Drosophila* tissues non-planar divisions are tolerated, with displaced cells reintegrating into the epithelial layer to preserve tissue integrity ^28, 46, 47^. Our live imaging analyses reveal that the time required for reintegration following non-planar divisions, a process also regulated by integrin function, is a critical, previously unrecognized factor in maintaining simple epithelial architecture. Notably, in egg chambers exhibiting multilayering, mutant cells within the ectopic layers show longer reintegration times compared to both control and mutant cells in non-multilayered egg chambers. Based on these observations, we propose that disruption of epithelial monolayer integrity concomitant with multilayer formation occurs only when reintegration time surpasses a defined threshold. Prolonged residence of displaced cells outside the epithelial layer may increase the likelihood of their subsequent division in the ectopic layer, thereby contributing to multilayering.

Our results further reveal that integrin mutant cells within ectopic layers exhibit higher surface tension than both control cells and mutant cells residing in monolayers. Consistent with previous findings that cells with elevated surface tension tend to segregate from surrounding populations ^34^, we observe that integrin mutant FCs progressively cluster over time. We propose that this increased cellular tension in integrin mutant cells within ectopic layers also contributes to multilayer formation. This is supported by our previous work showing that increasing cell tension enhances the multilayered phenotype associated to integrin loss of function ^48^. Moreover, this is consistent with *in vitro* studies of epidermal stratification, where cells that delaminate from the basal layer enter a high-tension state, an essential step believed to promote the transition from a single-layered epithelium to a stratified tissue by reinforcing the boundary between basal and suprabasal layers ^49^.

Altogether, we propose that the disruption of a simple epithelia and its transformation into a multilayered structure is governed by a multifactorial process, extending beyond the orientation of cell division alone (Fig. 6). Specifically, an increased proportion of cells dividing at non-parallel angles, combined with prolonged reintegration time for cells displaced into ectopic layers, impedes their return to the original monolayer and promotes the formation of additional layers. Moreover, elevated surface tension between cells within the ectopic layer further encourages their separation for the primary layer, reducing the likelihood of reintegration into a single-layered structure, thereby facilitating the stabilization and expansion of multilayering. Importantly, these processes are all regulated by integrins.

**Figure 6.**
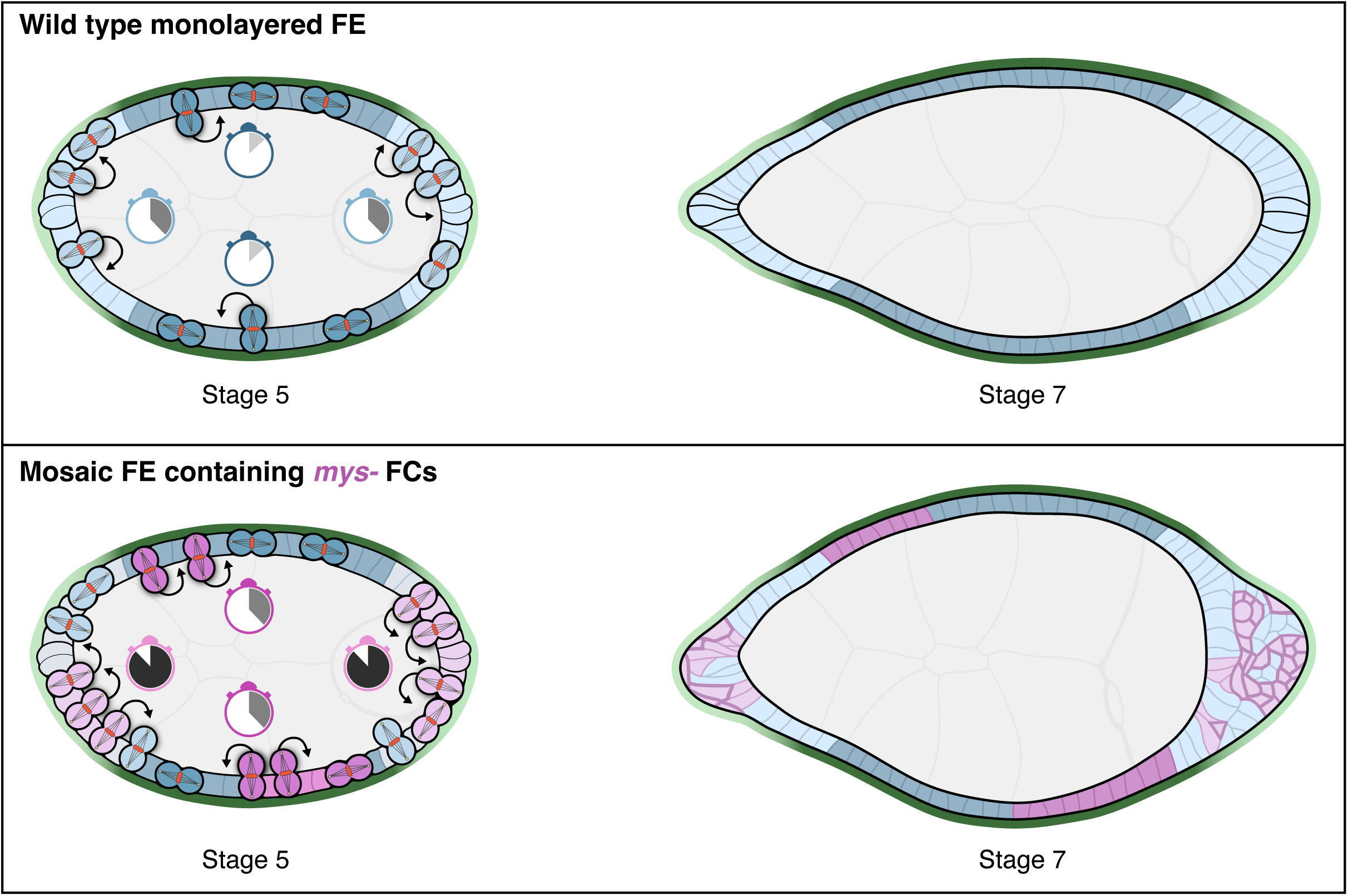
A model illustrating the mechanisms by which integrins and the mechanical properties of the BM align to safeguard simple epithelial architecture. Schematic of stage 5 or stage 7 wild type (top) and mosaic egg chambers containing *mys^-^* FCs (bottom). Anterior is left, posterior is right. The follicle cell monolayer is divided into terminal (light blue) and main body (dark blue) subdomains. Up to stage 6, FCs divide either parallel to the FE (planar divisions, both daughter cells stay in the monolayer) or at an angle (non-planar divisions, one daughter cell is displaced). In wild-type egg chambers (top), non-planar divisions are more frequent at the poles than in the mainbody, but epithelial architecture is maintained, as displaced cells reintegrate, with longer reintegration times at the poles (shown in chronometers). In mosaic egg chambers containing *mys^-^*FCs (bottom), non-planar divisions are more frequent, particularly at the poles. A gradient of BM stiffness, lower at the poles (green), along with prolonged reintegration times and increase contractility of *mys^-^* FCs in ectopic layers (stronger pink lines), contributes to the disruption of reintegration. This combination disturbs epithelial architecture and triggers multilayer formation.

Multilayering of the FE has been reported in mutants affecting different factors, including apico-basal polarity, spectrins and the Hippo tumour-suppressor pathway ^48, 50–52^. Intriguingly, in all cases, multilayering is restricted to the poles. The underlying reason for this spatial specificity in the disruption of simple architecture remains unclear. We propose that the mechanical properties of the BM, particularly its stiffness, play a crucial role in driving the localized formation of multilayering. Although existing evidence is limited and largely based on cell culture models, one study showed that a stiff BM significantly increases the proportion of planar-oriented divisions in resident skeletal muscle stem cells ^53^. Our findings extend this observation *in vivo* by revealing a correlation between BM stiffness and the orientation of the division plane within a developing simple epithelium. Specifically, we show that regions of the egg chamber with higher BM stiffness, such as the mainbody, exhibit an increased frequency of FC planar divisions. Furthermore, reintegration time following non-planar division inversely correlates with BM stiffness, with shorter reintegration times observed in the stiffer mainbody compared to the softer poles. Collectively, these results suggest that a stiff BM promotes planar divisions and rapid reintegration, thereby preserving monolayer integrity, whereas softer BM regions facilitate non-planar divisions and prolonged reintegration, predisposing these areas to epithelia disruption and multilayer formation, upon loss of factors regulating division orientation. However, our findings indicate that the loss of one of these factors, integrins, in an environment where the BM is uniformly soft does not result in multilayering. Based on this observation, we propose that in epithelial tissues, multilayer formation might not be driven solely by a soft BM, but rather by the juxtaposition of regions with differing BM stiffness, a mechanical feature not readily replicated in cell culture models. Such heterogeneity may facilitate localized transitions from single-layered to multilayered structures through the localized production of factors that soften the BM, a mechanism potentially critical for epithelia morphogenesis and repair. Supporting this notion, *in vivo* analysis of mammary gland branching in organotypic cultures have shown that duct elongation involves a reversible transition from single- or bilayered epithelia to a multilayered intermediate at the leading edge of the ducts, regions notably devoid of laminins ^54^.

Epithelial morphogenesis and tissue integrity are also influenced by geometric constraints, with epithelial cells in curved environments displaying distinct behaviors compared to those in flat contexts (reviewed in ^55^. For instance, in star-shaped epithelial monolayers, cell extrusion events are biased toward the tips, a pattern not observed in circular monolayers ^56^. Moreover, substrate curvature alone has been shown to promote stratification, growth and differentiation of corneal epithelial cells *in vitro* ^57^. During *Drosophila* oogenesis, the egg chamber initially spherical, undergoes anisotropic expansion along the anterior-posterior axis, elongating over two-fold, eventually forming an oval shape ^8, 16^. As a result, FCs at the poles experience greater curvature than those along the mainbody. Epithelial cells in such curved tissues can undergo intercalations along their apico-basal axis, adopting a shape known as “scutoid”, which facilitates cellular rearrangements ^58^. Remarkably, the degree of curvature correlates with the prevalence of scutoid formation, with more scutoids found at the poles of the egg chamber. Altogether, given that curved substrates facilitate cell rearrangement, extrusion and stratification, it is plausible that this geometrical constrains contribute to the observed multilayering at the poles. Moreover, tissue geometry can stem from the tissue’s intrinsic architecture, surrounding morphology, or BM mechanical properties (reviewed in ^55^. In the egg chamber, the BM stiffness is instructive for tissue elongation, with depletion of integrins, laminins, or ColIV disrupting elongation and resulting in rounder follicles ^8^. The fact that the uniform reduction of integrin, laminin or ColIV levels prevents multilayering, even in mosaic egg chambers containing *mys* clones, may stem not only from homogenization of the BM stiffness, but also from alterations in geometric constrains. Given the interdependence between egg chamber shape and BM mechanical properties, it is currently not possible to disentangle the independent contributions of BM stiffness and geometry in driving multilayering.

Tissue architecture emerges from the interplay between individual and collective cellular behaviors in response to molecular and mechanical cues. Our study reveals that integrins, together with BM mechanics are key to maintaining simple epithelial organization. Disruption of normal tissue architecture is a hallmark of oncogenic transformation. Forces from the underlying BM can affect epithelial tumor architecture and, in turn, modulate malignant potential ^59^. Identifying parameters regulating the maintenance of simple epithelia and their transition to stratified states is therefore not only essential for understanding epithelial morphogenesis, but also for elucidating mechanisms of cancer progression.

## Materials and methods

### Drosophila fly stocks and genetics

The following fly stocks were used: *mys*^11^ (also known as *mys^XG^*^43^ ^60^, *e22c-*Gal4-UAS-*flipase* ^61^, *tub* Gal80*^ts^* and w; *his:*RFP (Bloomington Drosophila Stock Centre, 7019 and 25377, respectively), UAS-*mys*RNAi (NIG-Fly, P{NIG.1560R}2), UAS-*LanB1* RNAi (VDRC, 23121), UAS-*ColIVα*2RNAi (VDRC, v106812), UAS-*asl*:YFP, *αtub:*GFP (a gift from Dr. Cayetano González), *resille*:GFP ^32^, *traffic jam*-Gal4 (*tj*-Gal4, ^39^).

The *e22c-*Gal4 driver is expressed in somatic cells of the germarium in the pupal and adult ovaries. It was used in combination with UAS-flipase and with *y w Ubi*-*GFP FRT-101* to generate *mys*^11^ clones. The heat shock flipase (*hs-flp*) system was also used to generate *mys*^11^ clones.

To observe *mys*^11^ clones produced after the germarium *mys^11^*FRT101/FMZ; *tub*Gal80*^ts^*/CyO females were crossed with *y w Ubi*-*GFP FRT-101; e22c-*Gal4 UAS*-flp* males. After eclosion, the middle of the flies was maintained 48 hours at 18°C as a control and the rest were 24 hours at 29°C as experimental.

To study wild type FC divisions *in vivo,* the stock *asl*:YFP,α*tub:*GFP; *his:*RFP was used. To study *mys*^11^ mutant FC divisions in monolayer *in vivo*, *asl*:YFP, *mys^11^*FRT19A/FMZ; *resille:*GFP-*his:*RFP females were crossed with *his*2AV:GFP*hs-flp*/FMZ; *e22c*-Gal4 UAS-*flp.* To study *in vivo mys*^11^ mutant FC divisions in multilayers and for laser ablation experiments, females *mys^11^*FRT101/FMZ; *resille*GFP were crossed with males *y w Ubi*-*GFP FRT-101; e22c-*Gal4 UAS*-flp.* In these experiments, flies were grown always at 25°C.

For rescue experiments *mys^11^*FRT19A/FMZ;; UAS-*LanB1*RNAi and *mys^11^*FRT19A/FMZ;; UAS-*ColIVα*2RNAi females were crossed with *hs-flp*RFPFRT19A/FMZ; *tj*-Gal4. In this case, flies were grown at 18°C and maintain at 25°C after eclosion.

### Ex vivo culturing of egg chambers

For live imaging, 1-2 days-old females were fattened on yeast during 2-3 days before dissection. We follow the protocol used by Villa-Fombuena *et al.*, 2021. Briefly, adult ovaries were dissected at room temperature in Ringer’s medium [128 mM NaCl, 2 mM KCl, 1.8 mM CaCl_2_, 4 mM MgCl_2_, 35.5 mM sucrose, 5 mM HEPES (pH 6.9)]. Ovarioles without the muscle sheath were transferred to a MatTek plate. The day before the experiment, we prepare in this plate a drop of 3 μl of Cell-Tak (Corning 354240) mixed with 3 μl of NaCOH_3_ and keep the plate at 4°C. We transfer the ovarioles into this drop. Then, the plate was supplemented with Schneider’s medium (Sigma Aldrich).

### Immunohistochemistry

1-2 days-old females were grown 2-3 days at 25°C before dissection. Adult ovaries were dissected at room temperature (RT) in Schneider’s medium (Sigma Aldrich), fixed in 4% paraformaldehyde in PBS (ChemCruz) for 20 min, permeabilized 30 min in PBT and blocked 1 h in PBT-10. Incubation with primary antibodies was performed overnight at 4°C in PBT-1. The following primary antibodies were used: chicken anti-GFP (1:600, Abcam Cat#13970), anti-GFP booster (1:500, Chromotek Cat#17303343), rat anti-RFP (1:500, Chromotek Cat#17812387). Secondary antibodies were incubated for 2 h in PBT-0.1 and used 1:200: anti-chicken Alexa Fluor 488 (Invitrogen Cat# A11039), TR anti-Rat (Sigma Aldrich Cat#SAB3700551). To label DNA, ovaries were incubated for 10 min with Hoechst (Sigma-Aldrich, 5 mg/ml; 1:1000 in PBT). For actin labelling, ovaries were incubated with Rhodamine Phalloidin (Biotium, 1:40) or 488 Phalloidin (Biotium, 1:40) during 15 minutes. Ovaries were mounted in Vectashield (Vector Laboratories).

PBT-10: PBS, 10% BSA, 1% tween20.

PBT-1: PBS, 1% BSA, 1% tween20.

PBT-0.1: PBS, 0.1% BSA, 1% tween20.

PBT: PBS (phosphate-buffered saline), 1% tween20.

### Laser ablation

Laser ablation experiments were performed in an Olympus IX-81 inverted microscope equipped with a spinning disk confocal unit (Yokogawa CSU-X1), a 100× oil objective, a 355 nm pulsed laser, third-harmonic, solid-state UV laser, and an Evolve 512 EMCCD digital camera (Photometrics). To analyze tension between FC-FC, a pulse of 125 mJ energy and 4 msec. duration, was applied to sever plasma membranes of cells. In all cases, cell surfaces were visualized with the membrane marker *Resille*-GFP and a Cobolt Calypso state laser (l = 491 nm 50 mW) was used for excitation of the GFP. Cuts between FC-FC were made parallel to the antero-posterior axis of the egg chamber. Images were taken 3 sec before and 10 sec after laser pulse, every 0,5 sec. The initial velocity was estimated as the velocity at the first time point (t1= 0,5 sec).

### Atomic force microscopy measurements

Atomic force microscopy measurements were performed as in ^35, 38^. Ovarioles were dissected out of the muscle sheath to make sure that the AFM cantilever was in direct contact with the BM. The cantilever was positioned at the desired position by brightfield microscopy. Statistical significance between experimental and control values was evaluated using a t-test, **** and *** P value < 0.00001 and <0.0001, respectively. Sixteen measurements were collected from each region (anterior, mainbody and posterior) across 6 control egg chambers, and 14 measurements per region from 6 *mys*-depleted egg chambers.

### Imaging and processing of samples

Live sample imaging was acquired on a Leica TCS-SP5 confocal microscope with 40x/1,3 oil objective and a Zeiss LSM 880 confocal microscope with a 40x/1,2 water. In all cases, intervals of 1 μm were taken and time points recorded every 1,5 minutes during 2-4 hours.

Fixed sample images were taken with Leica Stellaris confocal microscope with 40x/1,3 oil immersion objective. Z-stacks were 1 μm intervals.

Both cases, fixed and live images, were analyzed utilizing ImageJ and processed with Adobe Photoshop and Adobe Ilustrator.

### Statistical analysis

Statistical analysis of categorical data was done with *Chi square* test, comparing observed and expected frequencies (GraphPad QuickCalcs). To compare laser ablation measurements parametric *Student’s t* test was used. In the graphics the p-values of *Chi square* and *Student’s t* test are represented as: *= *P* value < 0.05, ** = *P* value < 0.01, *** = *P* value < 0.001 and ****= P value < 0,0001.

## Acknowledgements

We thank the BDSC for reagents. Research in our laboratories is funded by the Spanish Ministerio de Ciencia e Innovación and the FEDER programme (PID2019-109013GB-100 to M.D. M-B. and PIDXXX to A. G-R.) and by the Junta de Andalucía (Proyecto de Excelencia P20_00888).

## Author contributions

L.R-O, C.H. F-E designed research, performed experiments, analysed data and contributed to the writing of the manuscript. I.M.P performed experiments and provided useful comments on the manuscript. A. G-R provided financial support, designed research, analysed data and provided useful comments on the manuscript. M. D. M.-B. provided financial support, performed experiments, designed research, analysed data and wrote the manuscript.

## Competing interests

The authors declare no competing interests.

## Supplementary Information

**Supplementary Figure 1. Integrin mutant FCs undergoing planar division behave similarly to control.**

(A, B) Confocal images of live S6 egg chambers expressing histone-RFP (his:RFP, red) and histone-GFP (his:GFP, green), showing examples of planar divisions in the follicular epithelium of a control (A) and a mosaic egg chambers containing *mys* mutant follicle cell clones (B). Mutant FCs are identified by the absence of his:GFP (green). (C) Quantification of the frequency of control and mosaic egg chambers harbouring *mys* mutant clones that either remain in the FE (white in the columns) or exhibit reintegration times below 30 minutes (grey in the columns). Scale bar 3 μm (A, B).

**Supplementary Figure 2. Frequency and size of the multilayering increases with time.** (A-H) Control (A-D) and mosaic egg chambers containing *mys* mutant clones (E-H) at the indicated stages, stained with the nuclear marker Hoechst (white in A-H) and anti-GFP (green in E-H). Mutant FCs are identified by the absence of GFP signal. (I) Quantification of the multilayering phenotype at both anterior and posterior poles in mosaic egg chambers carrying *mys* mutant clones across the indicated stages. (J) Quantification of the frequency of mosaic egg chambers with *mys* mutant cells either interspersed with control cells or clustered together. (K, L) Mosaic egg chambers containing *mys* mutant clones at S5-6 (K) and 9-10 (L), stained with Hoechst (white in A-H) and anti-GFP (green in E-H). Mutant FCs lack GFP signal. (K’, L’) Magnified views of the regions outlined in white boxes in K and L, respectively. Scale bars 20 μm (A-H, K, L).

**Supplementary Figure 3. Downregulation of laminin or collagen IV levels rescue the multilayering phenotype cause by loss of integrin function in clones.**

(A-F) S9 egg chambers of the indicated genotypes stained with the nuclear marker Hoechst (white), anti-GFP (green) and the F-actin marker Rhodamine Phalloidin (RhPh, magenta). Mutant FCs are identified by the absence of GFP signal. (G) Quantification of the multilayering phenotype in egg chambers of the indicated genotypes. Scale bars 20 μm (A-F).

## Supplemental Movies

**Movie S1. Spindle orientation during planar division of a control FC located at the posterior pole.**

Time-lapse movie of a control FC located at the posterior pole undergoing planar division. Centrosomes are labelled with *asl*YFP (yellow), microtubules with *αtub*GFP (green) and DNA with *his*RFP (red). Scale bar, 3µm.

**Movie S2. Spindle orientation during non-planar division of a control FC located at the posterior pole.**

Time-lapse movie of a control FC located at the posterior pole undergoing non-planar division. Centrosomes are labelled with *asl*YFP (yellow), microtubules with *αtub*GFP (green) and DNA with *his*RFP (red). Scale bar, 3µm.

**Movie S3. Planar division of a control FC at the mainbody.**

Time-lapse movie of a control FC located at the mainbody undergoing planar division. DNA is labelled with *his2AV:GFP* and *his*RFP. Scale bar, 3µm.

**Movie S4. Planar division of a *mys* mutant FC at the posterior pole.**

Time-lapse movie of a *mys* mutant FC located at the posterior pole undergoing planar division. DNA is labelled with *his*RFP. Scale bar, 3µm.

**Movie S5. Non-planar division and reintegration of a control FC at the posterior pole.** Time-lapse movie of a control FC located at the posterior pole undergoing non-planar division. DNA is labelled with *his2AV:GFP* and *his*RFP. Scale bar, 3µm.

**Movie S6. Non-planar division and reintegration of a control FC at the mainbody.**

Time-lapse movie of a control FC located at the mainbody undergoing non-planar division. DNA is labelled with *his2AV:GFP* and *his*RFP. Scale bar, 3µm.

**Movie S7. Non-planar division and reintegration of a *mys* mutant FC at the mainbody.** Time-lapse movie of a *mys* mutant FC located at the mainbody undergoing non-planar division. DNA is labelled with *his*RFP. Scale bar, 3µm.

**Movie S8. Non-planar division and reintegration of a *mys* mutant FC at the posterior pole of a monolayered epithelium.**

Time-lapse movie of a *mys* mutant FC located at the posterior pole of a monolayered epithelium undergoing non-planar division. DNA is labelled with *his*RFP. Scale bar, 3µm.

**Movie S9. Non-planar division and reintegration of a *mys* mutant FC at the posterior pole of a multilayered epithelium.**

Time-lapse movie of a *mys* mutant FC located at the posterior pole of a multilayered epithelium undergoing non-planar division. DNA is labelled with *his*RFP. Scale bar, 3µm.

**Movie S10.**

Movie corresponds to the ablation experiment shown in Figure 4. The membranes of FCs are visualised with Resille-GFP. A cell bond between two control FCs is ablated. Scale bar: 3µm. GFP fluorescent is lost in the middle of the ablated bond upon laser ablation. The movie continues 10s after the cut and shows displacement of the vertexes. Images are taken every 0.5 seconds.

**Movie S11. Planar division of a FC at the posterior pole expressing a *mys*RNAi.**

Time-lapse movie of a FC expressing a *mysRNAi*, located at the posterior pole, undergoing planar division. Cell membrane is labelled with *Resille*GFP. Scale bar, 3µm.

**Movie S12. Non-planar division of a FC at the posterior pole expressing a *mys*RNAi.** Time-lapse movie of a FC expressing a *mysRNAi*, located at the posterior pole, undergoing non-planar division. Cell membrane is labelled with *Resille*GFP. Scale bar, 3µm.

## Notes

### Competing Interest Statement

The authors have declared no competing interest.

## References

1. Lechler, T. & Fuchs, E. Asymmetric cell divisions promote stratification and differentiation of mammalian skin. Nature 437, 275–280 (2005).

2. Smart, I.H. Variation in the plane of cell cleavage during the process of stratification in the mouse epidermis. Br J Dermatol 82, 276–282 (1970).

3. Williams, S.E., Beronja, S., Pasolli, H.A. & Fuchs, E. Asymmetric cell divisions promote Notch-dependent epidermal differentiation. Nature 470, 353–358 (2011).

4. Damen, M. et al. High proliferation and delamination during skin epidermal stratification. Nature communications 12, 3227 (2021).

5. Khalilgharibi, N. & Mao, Y. To form and function: on the role of basement membrane mechanics in tissue development, homeostasis and disease. Open Biol 11, 200360 (2021).

6. Agarwal, P., Shemesh, T. & Zaidel-Bar, R. Directed cell invasion and asymmetric adhesion drive tissue elongation and turning in C. elegans gonad morphogenesis. Dev Cell 57, 2111–2126 e2116 (2022).

7. Harmansa, S., Erlich, A., Eloy, C., Zurlo, G. & Lecuit, T. Growth anisotropy of the extracellular matrix shapes a developing organ. Nature communications 14, 1220 (2023).

8. Crest, J., Diz-Munoz, A., Chen, D.Y., Fletcher, D.A. & Bilder, D. Organ sculpting by patterned extracellular matrix stiffness. Elife 6 (2017).

9. Topfer, U., Guerra Santillan, K.Y., Fischer-Friedrich, E. & Dahmann, C. Distinct contributions of ECM proteins to basement membrane mechanical properties in Drosophila. Development 149 (2022).

10. Topfer, U. et al. AdamTS proteases control basement membrane heterogeneity and organ shape in Drosophila. Cell Rep 43, 114399 (2024).

11. Chen, J. & Krasnow, M.A. Integrin Beta 1 suppresses multilayering of a simple epithelium. PLoS One 7, e52886 (2012).

12. Fernández-Miñán, A., Martín-Bermudo, M.D. & Gonzéalez-reyes, A. Integrin signaling regulates spindle orientation in *Drosophila* to preserve the follicular-epithelium monolayer. Current Biology 17, 683–688 (2007).

13. Petridou, N.I. & Skourides, P.A. A ligand-independent integrin beta1 mechanosensory complex guides spindle orientation. Nature communications 7, 10899 (2016).

14. Bergstralh, D.T., Lovegrove, H.E. & St Johnston, D. Discs large links spindle orientation to apical-basal polarity in Drosophila epithelia. Curr Biol 23, 1707–1712 (2013).

15. Lovegrove, H.E., Bergstralh, D.T. & St Johnston, D. The role of integrins in Drosophila egg chamber morphogenesis. Development 146 (2019).

16. Spradling, A.C. Developmental genetics of oogenesis. The Development of Drosophila melanogaster. M. Bate and A Martinez-Arias, editors. Cold Spring Harbor Lab. Press, Cold Spring Harbor, *New York*, 1-70 (1993).

17. King, R.C. Ovarian development in *Drosophila* melanogaster. Academic Press, New York, NY. (1970).

18. Gutzeit, H.O., Eberhardt, W. & Gratwohl, E. Laminin and basement membrane-associated microfilaments in wild type and mutant *Drosophila* ovarian follicles. J. Cell Science 100, 781–788 (1991).

19. Calvi, B.R., Lilly, M.A. & Spradling, A.C. Cell cycle control of chorion gene amplification. Genes Dev 12, 734–744 (1998).

20. Gonzalez-Reyes, A. & St Johnston, D. Patterning of the follicle cell epithelium along the anterior-posterior axis during Drosophila oogenesis. Development 125, 2837–2846 (1998).

21. Haigo, S.L. & Bilder, D. Global tissue revolutions in a morphogenetic movement controlling elongation. Science 331, 1071–1074 (2011).

22. Goode, S. & Perrimon, N. Inhibition of patterned cell shape change and cel invasion by Discs large during *Drosophila* oogenesis. Genes and Development 11, 2532–2544 (1997).

23. Abdelilah-Seyfried, S., Cox, D.N. & Jan, Y.N. Bazooka is a permissive factor for the invasive behavior of *discs large* tumor cells in *Drosophila* ovarian follicular epithelia. Development 130, 1927–1935 (2003).

24. Bilder, D. Epithelial polarity and proliferation control: links from the Drosophila neoplastic tumor suppressors. Genes Dev 18, 1909–1925 (2004).

25. Lee, J.K., Brandin, E., Branton, D. & Goldstein, L.S. alpha-Spectrin is required for ovarian follicle monolayer integrity in Drosophila melanogaster. Development 124, 353–362 (1997).

26. Brown, N.H. Cell-cell adhesion via the ECM:integrin genetics in fly and worm. Matrix Biol. 19, 191–201 (2000).

27. Yee, G.H. & Hynes, R.O. A novel, tissue-specific integrin subunit, bg, expressed in the midgut of *Drosophila melanogaster*. Development 118, 845–858 (1993).

28. Bergstralh, D.T., Lovegrove, H.E. & St Johnston, D. Lateral adhesion drives reintegration of misplaced cells into epithelial monolayers. Nat Cell Biol 17, 1497–1503 (2015).

29. Jia, D., Xu, Q., Xie, Q., Mio, W. & Deng, W.M. Automatic stage identification of Drosophila egg chamber based on DAPI images. Sci Rep 6, 18850 (2016).

30. Santa-Cruz Mateos, C., Valencia-Exposito, A., Palacios, I.M. & Martin-Bermudo, M.D. Integrins regulate epithelial cell shape by controlling the architecture and mechanical properties of basal actomyosin networks. PLoS Genet 16, e1008717 (2020).

31. Farhadifar, R., Roper, J.C., Aigouy, B., Eaton, S. & Julicher, F. The influence of cell mechanics, cell-cell interactions, and proliferation on epithelial packing. Curr Biol 17, 2095–2104 (2007).

32. Morin, X., Daneman, R., Zavortink, M. & Chia, W. A protein trap strategy to detect GFP-tagged proteins expressed from their endogenous loci in Drosophila. Proc Natl Acad Sci U S A 98, 15050–15055 (2001).

33. Hutson, M.S. et al. Forces for morphogenesis investigated with laser microsurgery and quantitative modeling. Science 300, 145–149 (2003).

34. Brodland, G.W. The Differential Interfacial Tension Hypothesis (DITH): a comprehensive theory for the self-rearrangement of embryonic cells and tissues. J Biomech Eng 124, 188–197 (2002).

35. Diaz de la Loza, M.C., et al. Laminin Levels Regulate Tissue Migration and Anterior-Posterior Polarity during Egg Morphogenesis in Drosophila. Cell Rep 20, 211–223 (2017).

36. Martin-Bermudo, M.D. & Brown, N.H. Uncoupling integrin adhesion and signaling: the b_PS_ cytoplasmic domain is sufficient to regulate gene expression in the *Drosophila* embryo. Genes Dev. 13, 729–739 (1999).

37. Valencia-Exposito, A., Gomez-Lamarca, M.J., Widmann, T.J. & Martin-Bermudo, M.D. Integrins Cooperate With the EGFR/Ras Pathway to Preserve Epithelia Survival and Architecture in Development and Oncogenesis. Front Cell Dev Biol 10, 892691 (2022).

38. Molina Lopez, E., et al. Constriction imposed by basement membrane regulates developmental cell migration. PLoS Biol 21, e3002172 (2023).

39. Li, M.A., Alls, J.D., Avancini, R.M., Koo, K. & Godt, D. The large Maf factor Traffic Jam controls gonad morphogenesis in Drosophila. Nat Cell Biol 5, 994–1000 (2003).

40. Jones, P.H. & Watt, F.M. Separation of human epidermal stem cells from transit amplifying cells on the basis of differences in integrin function and expression. Cell 73, 713–724 (1993).

41. Simpson, C.L., Patel, D.M. & Green, K.J. Deconstructing the skin: cytoarchitectural determinants of epidermal morphogenesis. Nat Rev Mol Cell Biol 12, 565–580 (2011).

42. Brakebusch, C. et al. Skin and hair follicle integrity is crucially dependent on beta 1 integrin expression on keratinocytes. EMBO J 19, 3990–4003 (2000).

43. Byrd, K.M. et al. LGN plays distinct roles in oral epithelial stratification, filiform papilla morphogenesis and hair follicle development. Development 143, 2803–2817 (2016).

44. Schaeffer, V., Althauser, C., Shcherbata, H.R., Deng, W.M. & Ruohola-Baker, H. Notch-dependent Fizzy-related/Hec1/Cdh1 expression is required for the mitotic-to-endocycle transition in Drosophila follicle cells. Curr Biol 14, 630–636 (2004).

45. Seldin, L., Muroyama, A. & Lechler, T. NuMA-microtubule interactions are critical for spindle orientation and the morphogenesis of diverse epidermal structures. Elife 5 (2016).

46. Ciruna, B., Jenny, A., Lee, D., Mlodzik, M. & Schier, A.F. Planar cell polarity signalling couples cell division and morphogenesis during neurulation. Nature 439, 220–224 (2006).

47. Packard, A. et al. Luminal mitosis drives epithelial cell dispersal within the branching ureteric bud. Dev Cell 27, 319–330 (2013).

48. Ng, B.F. et al. alpha-Spectrin and integrins act together to regulate actomyosin and columnarization, and to maintain a monolayered follicular epithelium. Development 143, 1388–1399 (2016).

49. Miroshnikova, Y.A. et al. Adhesion forces and cortical tension couple cell proliferation and differentiation to drive epidermal stratification. Nat Cell Biol 20, 69–80 (2018).

50. Tanentzapf, G., Smith, C., McGlade, J. & Tepass, U. Apical, Lateral, and Basal polarization cues contribute to the development of the follicular epithelium during *drosophila* oogenesis. *J*. Cell Biology 151, 891–904 (2000).

51. Meignin, C., Alvarez-Garcia, I., Davis, I. & Palacios, I.M. The salvador-warts-hippo pathway is required for epithelial proliferation and axis specification in Drosophila. Curr Biol 17, 1871–1878 (2007).

52. Polesello, C. & Tapon, N. Salvador-warts-hippo signaling promotes Drosophila posterior follicle cell maturation downstream of notch. Curr Biol 17, 1864–1870 (2007).

53. Moyle, L.A. et al. Three-dimensional niche stiffness synergizes with Wnt7a to modulate the extent of satellite cell symmetric self-renewal divisions. Mol Biol Cell 31, 1703–1713 (2020).

54. Ewald, A.J., Brenot, A., Duong, M., Chan, B.S. & Werb, Z. Collective epithelial migration and cell rearrangements drive mammary branching morphogenesis. Dev Cell 14, 570–581 (2008).

55. Nelson, C.M. Geometric control of tissue morphogenesis. Biochim Biophys Acta 1793, 903–910 (2009).

56. Saw, T.B. et al. Topological defects in epithelia govern cell death and extrusion. Nature 544, 212–216 (2017).

57. Gouveia, R.M., Koudouna, E., Jester, J., Figueiredo, F. & Connon, C.J. Template Curvature Influences Cell Alignment to Create Improved Human Corneal Tissue Equivalents. Adv Biosyst 1, e1700135 (2017).

58. Gomez-Galvez, P. et al. Scutoids are a geometrical solution to three-dimensional packing of epithelia. Nature communications 9, 2960 (2018).

59. Fiore, V.F. et al. Mechanics of a multilayer epithelium instruct tumour architecture and function. Nature 585, 433–439 (2020).

60. Bunch, T.A. & Brower, D.L. Drosophila PS2 integrin mediates RGD-dependent cell-matrix interactions. Development 116, 239–247 (1992).

61. Duffy, J.B., Harrison, D.A. & Perrimon, N. Identifying loci required for follicular patterning using directed mosaics. Developmen 125, 2263–2271 (1998).

